# Romer-EPTI: rotating-view motion-robust super-resolution EPTI for SNR-efficient distortion-free in-vivo mesoscale dMRI and microstructure imaging

**DOI:** 10.1101/2024.01.26.577343

**Authors:** Zijing Dong, Timothy G. Reese, Hong-Hsi Lee, Susie Y. Huang, Jonathan R. Polimeni, Lawrence L. Wald, Fuyixue Wang

## Abstract

**Purpose:** To overcome the major challenges in dMRI acquisition, including low SNR, distortion/blurring, and motion vulnerability.

**Methods:** A novel Romer-EPTI technique is developed to provide distortion-free dMRI with significant SNR gain, high motion-robustness, sharp spatial resolution, and simultaneous multi-TE imaging. It introduces a ROtating-view Motion-robust supEr-Resolution technique (Romer) combined with a distortion/blurring-free EPTI encoding. Romer enhances SNR by a simultaneous multi-thick-slice acquisition with rotating-view encoding, while providing high motion-robustness through a motion-aware super-resolution reconstruction, which also incorporates slice-profile and real-value diffusion, to resolve high-isotropic-resolution volumes. The in-plane encoding is performed using distortion/blurring-free EPTI, which further improves effective spatial resolution and motion robustness by preventing not only T_2_/T_2_*-blurring but also additional blurring resulting from combining encoded volumes with inconsistent geometries caused by dynamic distortions. Self-navigation was incorporated to enable efficient phase correction. Additional developments include strategies to address slab-boundary artifacts, achieve minimal TE for SNR gain at 7T, and achieve high robustness to strong phase variations at high b-values.

**Results:** Using Romer-EPTI, we demonstrate distortion-free whole-brain mesoscale in-vivo dMRI at both 3T (500-μm-iso) and 7T (485-μm-iso) for the first time, with high SNR efficiency (e.g., 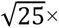), and high image quality free from distortion and slab-boundary artifacts with minimal blurring. Motion experiments demonstrate Romer-EPTI’s high motion-robustness and ability to recover sharp images in the presence of motion. Romer-EPTI also demonstrates significant SNR gain and robustness in high b-value (b=5000s/mm^2^) and time-dependent dMRI.

**Conclusion:** Romer-EPTI significantly improves SNR, motion-robustness, and image quality, providing a highly efficient acquisition for high-resolution dMRI and microstructure imaging.

## 1. Introduction

The evolution in in-vivo diffusion MRI (dMRI) has yielded exciting insights into the structural connectivity and tissue microstructure of the human brain. Notably, high-resolution dMRI techniques (1–8) have enabled the investigation of the brain’s fine-scale structures, allowing for the visualization of small but crucial circuitries (1,3,9,10) and the cytoarchitecture within the thin cortical layers (2,5). Moreover, hardware advances (11–14) have made it possible to use stronger diffusion encodings/gradients (15–20). Combined with advanced diffusion models (21–30), high-b value dMRI can probe various tissue microstructural characteristics with more comprehensive and precise information. Advancing dMRI in these directions holds great promise of providing richer information about the human brain structure in both health and disease (31–33). However, significant technical challenges persist in in-vivo dMRI acquisition that limit its ability to achieve higher spatial resolution and/or increased b-values. These challenges include: i) diminishing signal-to-noise ratio (SNR) as voxel size is reduced and/or b-values are raised; ii) severe image degradation, including geometric distortion and T_2_/T_2_*-blurring; and iii) susceptibility to subject motion and motion-related phase variations. Over the past decades, significant efforts have been made to address these limitations in in-vivo dMRI.

To improve SNR efficiency, several acquisition methods have been developed. Notably, multi-slab imaging (4,6,34–36) employs thick-slab acquisition with Fourier encodings along slice direction, and reconstructs thin-slice volume from these encoded thick-slab data acquired across multiple TRs. The thick-slab acquisition enhances SNR efficiency by reducing TR and achieving more efficient noise averaging. Simultaneous multi-slab imaging (37–39) combines multi-slab imaging with simultaneous multi-slice (SMS) (40–42) to further improve the SNR efficiency. Similarly, gSlider-SMS (7,43,44) employs thick-slab acquisition with Hadamard-like encodings along slice direction (45,46) for self-navigated phase correction, and combines with SMS for higher SNR efficiency. However, despite the efficacy of these methods in improving SNR, they suffer from several limitations. Both the multi-slab imaging and gSlider suffer from slab boundary artifacts, manifesting as striping artifacts at slab boundaries due to imperfect slab profiles and/or spin-history effects. This issue has been recognized and several methods have been developed to mitigate these artifacts (44,47,48). Furthermore, to estimate and correct physiological-motion-induced phase variations between encoded volumes, these methods often require additional navigator acquisition (34), or specialized RF pulses (7) for self-navigated spatial encoding, leading to either compromised efficiency with increased specific absorption rate (SAR) deposition or susceptibility to *B*_0_/*B*_1_ inhomogeneity, which can pose challenges at higher field strengths. Alternatively, another promising strategy, the super-resolution methods (49–55) can be used to improve SNR efficiency by acquiring multiple low-resolution volumes with different slice shifts, orthogonal views, or orientations to resolve a high-resolution volume. They provide a viable approach for self-navigated phase correction without the cost of additional scan time, extra SAR deposition, or *B*_0_/*B*_1_ susceptibility. However, all aforementioned methods share two additional common drawbacks. First, they suffer from distortion and T_2_*-blurring artifacts, particularly severe at high spatial resolutions and/or high field strengths. Second, they can be sensitive to subject motion. In addition to the physiological-motion-induced phase variations as mentioned above, several effects associated with the bulk motion between spatially-encoded volumes can interrupt spatial encoding and lead to motion artifacts. While it is feasible to model bulk motion in the reconstruction process (43), it can be challenging to fully address motion-related artifacts—notably the motion-related dynamic distortions that can be difficult to model. Motion can change the susceptibility field, the direction of the PE axis, and the interaction between the head position and the eddy-current or the shimming fields, leading to changes in image geometry. As a result of combining encoded volumes with inconsistent geometries/dynamic distortions, detrimental artifacts and blurring will be introduced on the final reconstructed high-resolution images. All these limitations (striping artifacts, SAR, *B*_0_/*B*_1_ susceptibility, distortion, blurring, motion) limit the attainable image quality and effective spatial resolution, and pose escalating challenges at higher resolutions, b-values, and field strengths.

To reduce distortion and blurring of EPI, parallel imaging (56–58) and multi-shot EPI including readout-segmented (59–63) and interleaved-segmented methods (64–73), can be used. However, achieving more effective distortion/blurring reduction requires a higher acceleration factor or a larger number of shots, resulting in higher noise amplification or prolonged scan time, and residual distortions still remain. Methods using reversed phase-encoding acquisitions to derive a field map, and subsequently correcting distortion either through post-processing (e.g., TOPUP (74,75)) or integration into reconstruction (e.g., BUDA (76,77)) can correct for image distortion but do not alleviate image blurring. To fully eliminate distortion and blurring, a point-spread-function (PSF) encoded multi-shot EPI approach was developed (78–81). Our tilted-CAIPI technique (82) further accelerates PSF-EPI from ∼200 shots per image (fully-sampled) to less than 10 shots at 1-mm resolution, achieving distortion/blurring-free dMRI within a practical scan time. More recently, we developed Echo-planar time-resolving imaging (EPTI) (83–89) to achieve distortion/blurring-free time-resolved multi-echo imaging. Instead of combing PE lines with different phase accumulation and signal decay to generate a single distorted and blurred image as in EPI, EPTI reconstructs the *k*_y_-*t* (PE-TE) space data and resolves a time series of distortion/blurring-free images within the readout at a short TE interval of ∼1 ms. Moreover, EPTI can provide additional SNR gain over EPI (30-40%) for diffusion imaging by achieving minimized TE using echo-train shifting and optimized readout lengths (86).

In this work, to overcome the challenges in dMRI acquisition, we introduce a novel technique, ROtating-view Motion-robust supEr-Resolution (Romer) EPTI, which can achieve: **i**) significantly higher SNR efficiency, for example, a 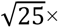 folds improvement compared to conventional EPI, corresponding to a 25× scan time reduction; **ii)** high-quality images free from distortion and slab boundary striping artifacts, with minimal spatial blurring; **iii)** high robustness to various source of motion (3D bulk motion, physiological motion, and motion-related dynamic distortions); **iv**) minimal TE, no extra SAR or *B*_0_/*B*_1_ susceptibility for 7T dMRI; **v**) simultaneously resolved multi-TE images for diffusion-relaxometry.

Romer-EPTI introduces an SNR-efficient rotating-view motion-robust super-resolution technique (Romer), which encodes along slice-readout dimensions, combined with a distortion/blurring-free self-navigated EPTI encoding, which encodes along the PE-TE dimensions. Specifically, Romer enhances SNR by a simultaneous multi-thick-slice acquisition with rotating-view encoding, while ensuring high motion robustness through a motion-aware super-resolution reconstruction, which also incorporates real-value diffusion and realistic slice profile, to resolve high-isotropic-resolution volumes. The in-plane encoding is performed using distortion/blurring-free EPTI readout, which further improves effective spatial resolution and motion robustness as it prevents not only T_2_/T_2_* blurring but also additional blurring/artifacts in the final high-resolution image that result from the combination of encoded volumes with inconsistent geometries caused by dynamic distortion. Romer-EPTI encoding is developed with self-navigation capability to enable effective removal of physiological-motion-related phase variations without extra scan time, SAR, or increased *B*_0_/*B*_1_ susceptibility. Additional developments have been incorporated into Romer-EPTI to further improve its performance, including an acquisition scheme to address slab-boundary artifacts, an optimized EPTI readout to achieve minimal TE for additional SNR gain, and single-shot EPTI encoding to achieve high robustness to strong phase variations at high-b values.

Using Romer-EPTI, we demonstrate distortion-free whole-brain in-vivo dMRI acquired at a mesoscopic resolution (∼0.1 mm^3^, ∼27× smaller voxel than the standard 1.5-mm isotropic resolution) at both 3T (500-μm isotropic) and 7T (485-μm isotropic) on clinical scanners. Initial results of this work have been presented at the ISMRM (90–92). To our knowledge, this is the first study that has achieved whole-brain mesoscale in-vivo dMRI on both 3T and 7T. Motion experiments demonstrate the high motion robustness of Romer-EPTI and its ability to recover high-quality sharp diffusion images in the presence of subject motion. We also demonstrate its significant SNR gain and robustness for microstructure imaging, such as high b-value dMRI (e.g., at 5000 s/mm^2^) and time-dependent diffusion.

## 2. Theory

### 2.1 Romer: ROtating-view Motion-robust supEr-Resolution

#### Thick-slice acquisition with rotating-view encoding

As shown in Fig. 1, Romer acquires thick-slice volumes with reduced volume TR to enhance SNR efficiency. Multi-band excitation with blipped CAIPI encoding (40–42) can be applied to acquire multiple thick slices simultaneously to further shorten the volume TR and improve SNR efficiency. Then, a rotating-view super-resolution scheme is employed across different thick-slice volumes to encode and recover the high isotropic resolution. For instance, by rotating the slice-readout direction around the PE axis, Romer samples thick-slice volumes with different slice orientations. The rotating-view encoding is designed to sample sufficient spatial information to achieve full reconstruction of the target spatial resolution. For example, if we define the ratio of the slice thickness between the encoded thick-slice volume and the target thin-slice volume as *Romer_factor_*, the number of encoding orientations (*N*) should be no less than π/2 times 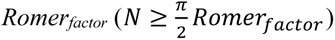 (51,93). From a *k*-space sampling perspective, the thick-slice sampling in Romer can be approximated as a blade-band encoding in the 3D *k*-space (with low-frequency along *k*_z_ and high-frequency along *k*_x_). The rotation of slice orientation corresponds to the rotation of the blades in the 3D *k*-space. By acquiring rotated blades with multiple orientations fulfilling the Nyquist criterion, the entire 3D *k*-space is sampled including the high-frequency signals needed to produce an isotropic high-resolution volume.

**Figure 1.**
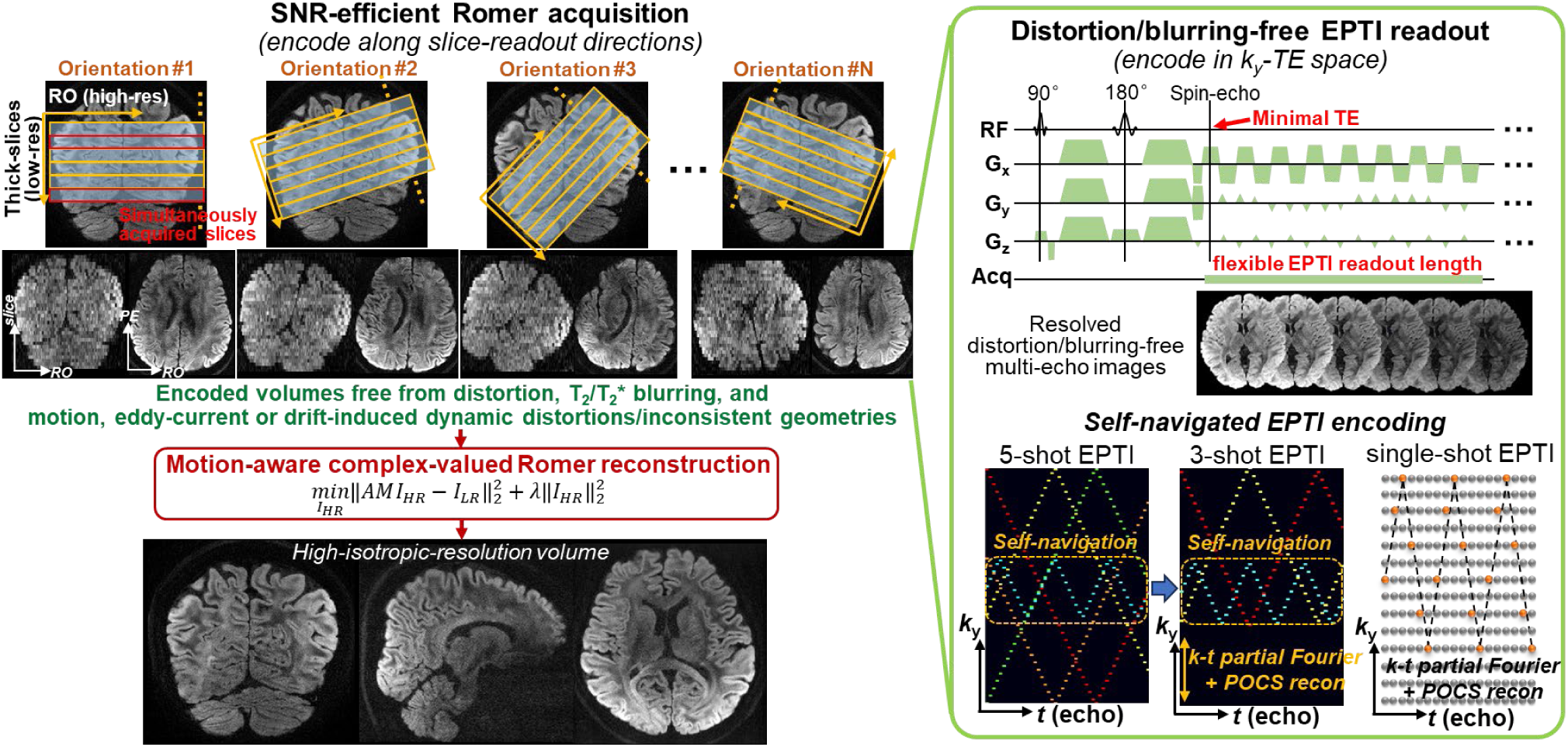
Illustration of Romer-EPTI. The Romer acquisition enhances SNR efficiency with high motion robustness. It encodes the slice-readout dimensions by acquiring thick-slice volumes with different slice orientations. Multi-band excitation can be applied to acquire multiple thick slices simultaneously to further improve SNR efficiency. The in-plane encoding of the Romer-encoded volumes is performed using EPTI readout, so these encoded volumes are free from i) geometric distortion, ii) T_2_/T_2_* blurring, and iii) motion, eddy-current or drift-induced dynamic distortions/geometric inconsistencies between the encoded thick-slice volumes, which can cause detrimental artifacts/blurring on the final reconstructed high-resolution image. Then, a motion-aware super-resolution reconstruction that incorporates 3D bulk motion, realistic slice profile, and real-value diffusion, is proposed to reconstruct the target high-isotropic-resolution volume. The sequence diagram of the diffusion EPTI readout and the self-navigated EPTI encodings are presented on the right. EPTI encodes the *k*_y_-TE space and resolves multi-echo distortion/blurring-free images across the readout. The continuous EPTI readout enables a flexible readout length that is independent of spatial matrix size and can be chosen to achieve optimized SNR efficiency. The TE can be minimized by starting the readout immediately at the spin-echo point to avoid deadtime between 90° and 180° pulses to further improve SNR. For mesoscale imaging, a 5-shot EPTI encoding is designed and can be further accelerated to 3-shot using a *k-t* partial Fourier acquisition. For high-b value imaging, a single-shot EPTI encoding is employed to provide high robustness to physiological phase variations.

In addition, to minimize potential spin-history artifacts associated with a short TR, we use an acquisition ordering scheme that acquires diffusion encodings or averages first before orientations. This sampling order avoids spatially non-uniform signal recovery that may occur when changing orientations for each volume. Furthermore, since there is no accumulative spin-history effect at the edge of the thick slices across different Romer-encoded volumes, the final reconstructed high-resolution volumes will be free from the slab-boundary striping artifacts.

#### Motion-aware super-resolution reconstruction

In Romer, various sources of motion-related effects between the encoded thick-slice volumes are addressed to ensure the reconstruction of high-fidelity volumes with sharp effective spatial resolution. These include (a) 3D bulk motion; (b) physiological-motion induced phase variations; and (c) geometric inconsistencies/dynamic distortions due to motion, eddy-current, or drift-induced field changes.

The rotating-view encoding employed in Romer itself is highly suitable for motion correction (effect a) because of its oversampled center with data redundancy. To correct for 3D bulk motion, a motion-aware super-resolution reconstruction is employed. The following minimization problem is solved to obtain each high-resolution volume:

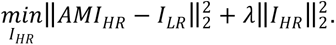

Here, *I_LR_* represents the thick-slice volumes acquired with rotated slice orientations, *I_HR_* is the target high-resolution volume, *M* is the rigid-motion transformation matrix (94) modeling 3D motion between the encoded thick-slice volumes, *A* is the Romer encoding matrix constructed based on the slice profile calculated from the 90°-180° RF pulses, and λ controls the Tikhonov regularization. The Romer acquisition provides self-navigation capability as each thick-slice volume can be reconstructed independently with high SNR, and the motion parameters can be directly estimated by registering the thick-slice volumes after up-sampling them to the same isotropic resolution.

To address the physiological motion-induced phase variations (effect b), the spatially low-frequency 3D phases are removed from each Romer-encoded volume in a self-navigation manner. Additionally, to address geometry changes/dynamic distortions (effect c), the in-plane encoding of the Romer-encoded thick-slice volumes is performed using a distortion-free EPTI readout, as described in section 2.2.

#### Complex-valued reconstruction and slice-profile modeling

The proposed Romer reconstruction is designed to perform complex-valued reconstruction after removing the low-frequency phase differences between the thick-slice volumes. This allows the use of real-value data (95) rather than magnitude data, thereby reducing the noise floor (Supporting Information Figure S1A). Additionally, Romer incorporates realistic slice profiles in the reconstruction model. Although Romer is relatively robust to slice profile variations as it does not rely on RF-encoding to resolve spatial resolution, neglecting the imperfect slice profile (e.g., using simplified model like box function) can still introduce bias. By using realistic slice profiles that account for the modulation within the slice and the cross-talk into the neighboring slices, we can further improve the reconstruction accuracy and prevent information loss (Supporting Information Figure S1B).

### 2.2 Romer with EPTI encoding

The in-plane encoding of the Romer-encoded volumes is performed using EPTI readout. This not only eliminates distortion and T_2_/T * blurring (e.g., 45% PSF side-lobe amplitude in EPI even with a 4× in-plane acceleration at 500 μm, Fig. 2A-a) and achieves sharp effective spatial resolution along the phase-encoding direction, but also further improves the motion robustness of Romer. Specifically, it eliminates geometric inconsistencies (i.e., dynamic distortions) between the encoded thick-slice volumes caused by motion, eddy-current, or drift-induced field changes, therefore avoiding additional detrimental artifacts/blurring on the final reconstructed high-resolution images due to the combination of these inconsistent geometries. For example, at 500-μm resolution, EPI readout even with a 4× in-plane acceleration can result in ∼400% blurring due to dynamic distortions (Fig. 2A-b), while EPTI provides consistent geometry of the spatially-encoded volumes and eliminates these sources of blurring (Fig. 2A-c).

**Figure 2.**
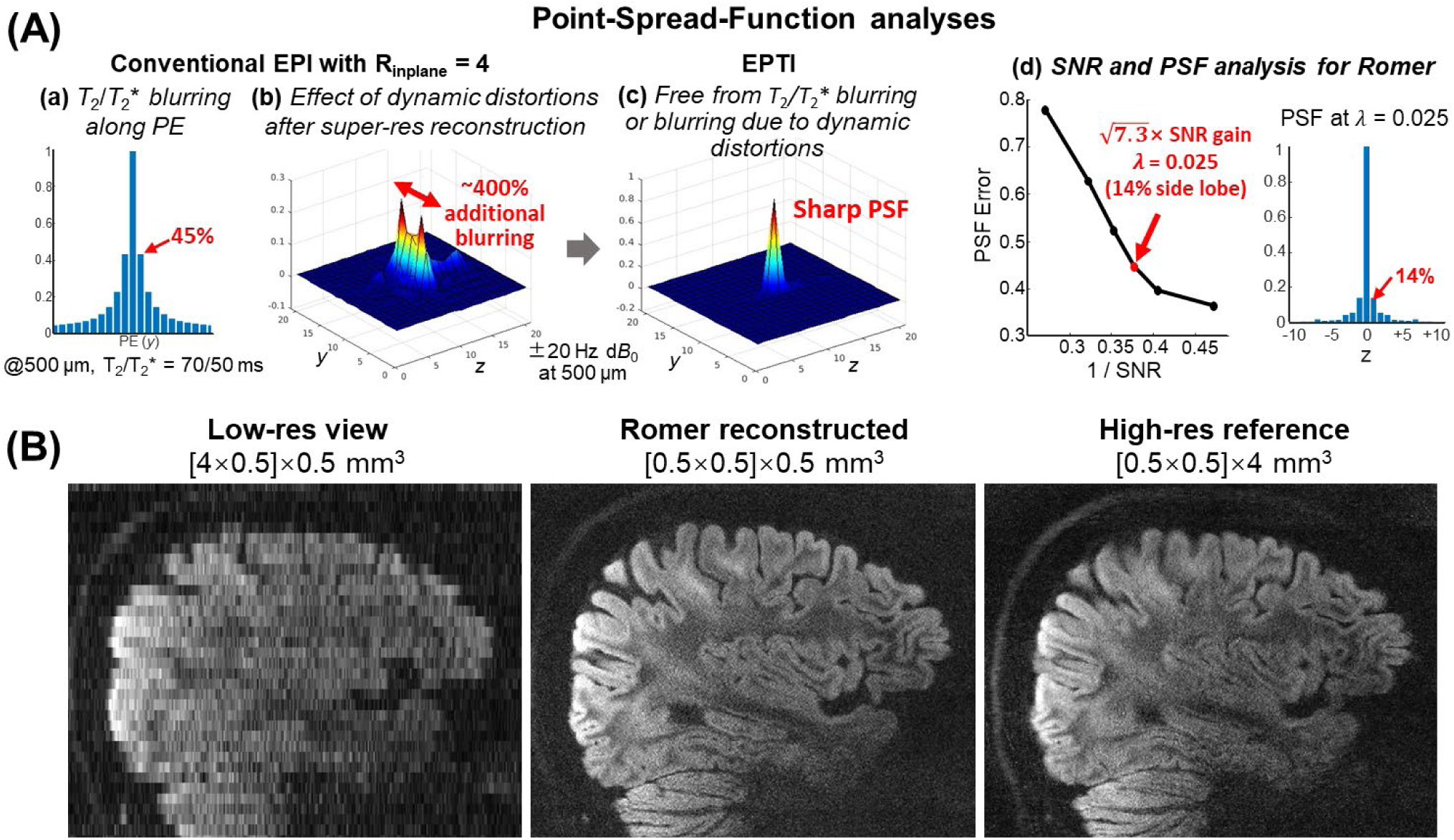
(**A**) Point-spread-function (PSF) analyses. a) PSF that shows the T_2_/T_2_* blurring of EPI along the phase encoding (PE) direction due to signal decay across readout, with a 45% side-lobe amplitude at 500-μm resolution even with a high in-plane acceleration of 4. b) PSF that shows additional image blurring on the final reconstructed high-resolution image using EPI readout, which results from combining encoded volumes with dynamic distortion/inconsistent geometries caused by motion, eddy-current, or drift induced field changes. A ±20 Hz *dB*_0_ change can lead to ∼400% wider PSF at 500-μm using EPI even with a high in-plane acceleration factor of 4, significantly compromising the effective spatial resolution. c) PSF of the final reconstructed high-resolution volume using EPTI readout to perform in-plane encoding. The sources of blurring (T_2_/T_2_* blurring and blurring due to dynamic distortions) are eliminated and a sharp PSF is achieved. d) PSF-error (along slice-readout directions) versus 1/SNR (L-curve) under different Tikhonov regularization parameters in Romer reconstruction. At a good trade-off point between PSF error and SNR (λ = 0.025), a Romer factor of 8 can achieve a high SNR efficiency gain of 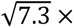 (close to the theoretical SNR gain of √8 ×), while achieving sharp resolution along slice-readout dimensions with a mild spatial blurring (∼14% side-lobe amplitude) at a mesoscale resolution of 500 μm. This, combined with SMS and optimized EPTI readout, achieves an overall SNR efficiency gain of 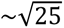 folds. (**B**) Comparison of the low-resolution view of a thick-slice volume (left), the Romer reconstructed high-isotopic-resolution volume at the same view (middle), and the reference high in-plane resolution EPTI image at the same view (right). Romer effectively reconstructs the isotropic resolution volume with sharp resolution and no noticeable blurring compared to the directly sampled reference high in-plane resolution data.

EPTI exploits the spatiotemporal correlation to reconstruct the full *k*_y_-*t/TE* space and resolve a time series of multi-echo images at every echo spacing (Fig. 1, right). Because of this time-resolving feature, EPTI enables continuous sampling with highly flexible readout length and TEs. Specifically, EPTI’s readout length is independent of spatial matrix sizes, and can be optimized to improve SNR efficiency. In addition, it allows for flexible TEs without constraints imposed by readout length as in EPI, and a minimal TE can be achieved by starting the readout immediately at the spin-echo timepoint, thereby reducing deadtime between 90° and 180° pulses. Together, they provide additional SNR gain on top of the Romer acquisition. The multi-TE images resolved across EPTI readout also provide useful information for diffusion-relaxometry and multi-compartment modeling (25,96–98).

The EPTI encoding is designed to sample *k*-space center in each shot (Fig. 1, bottom right) to provide self-navigation for physiological phase correction (86). To achieve mesoscale in-plane resolution, a 5-shot EPTI encoding is designed, and can be accelerated to 3 shots using a *k-t* partial Fourier acquisition (with POCS (99) to preserve spatial resolution). To further improve the robustness of Romer-EPTI to phase variations for high/ultra-high b-value imaging, a single-shot EPTI encoding (92,100,101) is employed, which has shown robust performance at standard spatial resolutions (e.g., 1.2–3 mm). Subspace reconstruction (84,102–104) with shot-to-shot phase correction (86) is used to reconstruct EPTI data.

#### SNR efficiency provided by Romer-EPTI

The overall SNR improvement provided by Romer-EPTI is determined by *Romer_factor_*, multiband (MB) factor, EPTI readout, and Romer reconstruction. We characterize the SNR gain of Romer through an SNR and PSF analysis, which considers the Romer reconstruction conditioning and the effect of regularization, similar to the approach described in Wang et al (43). A higher Tikhonov regularization improves the conditioning and leads to higher SNR, but will result in higher PSF error—a measure of effective resolution. Fig. 2A-d shows that, at a good trade-off point between SNR and PSF error, a Romer factor of 8 can achieve a high SNR efficiency gain of 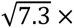 (close to the theoretical SNR gain of √8 ×), while achieving sharp resolution along slice-readout dimensions with mild spatial blurring (∼14% side-lobe amplitude). By combining with an MB factor of 2, Romer provides a 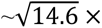 SNR efficiency gain 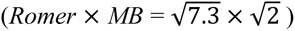. Then, as mentioned above, the EPTI readout with minimized TE and optimized readout length contributes an additional ∼30-40% SNR gain (86), leading to an overall SNR efficiency gain of 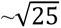 folds 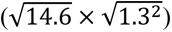, compared to conventional 2D slice-by-slice imaging as a benchmark.

## 3. Methods

In-vivo experiments were performed on healthy volunteers with a consented institutionally approved protocol on a Siemens Prisma 3T scanner with a 32-channel head receiver coil (Siemens Healthineers, Erlangen, Germany), and on a Siemens Terra 7T scanner with a 32-channel Nova head coil (Nova Medical, Wilmington, MA).

### 3.1 In-vivo mesoscale dMRI at 500 μm-iso resolution at 3T

Whole-brain mesoscale in-vivo diffusion data were acquired at 500-μm isotropic resolution on a 3T Prisma using Romer-EPTI to demonstrate its SNR efficiency and image quality. The data were acquired with imaging parameters: b = 1000 s/mm^2^, 25 diffusion directions, FOV = 200×192×184 mm^3^, matrix size of the high-resolution volume = 400×384×368, matrix size of the acquired thick-slice volume = 400×384×46, *Romer_factor_* of 8, 12 orientations (15° rotation per orientation), 5-shot EPTI encoding, MB factor = 2, 56 echoes, TE_range_ = 39–123 ms, echo spacing = 1.54 ms, TR = 3.1 s, total acquisition time ∼80 minutes. As a comparison, thick-slice EPI data was acquired with the following parameters: FOV = 200×192×184 mm^3^, resolution = 0.5×0.5×4 mm^3^, MB factor = 2, in-plane acceleration factor (R_inplane_) = 4, partial Fourier factor = 5/8, TE/TR = 77/3700 ms. Even after using a high acceleration (R_inplane_ = 4) and a partial Fourier factor of 5/8, EPI has a longer TE than EPTI (77 ms vs 39 ms) and a long TR due to the prolonged echo train length.

### 3.2 In-vivo motion experiment at 500 μm-iso resolution

To evaluate the motion robustness of Romer-EPTI, an in-vivo motion experiment was conducted at 500-μm isotropic resolution. The identical 3T mesoscale Romer-EPTI imaging protocol was used with a 3-shot EPTI encoding with a *k-t* Partial Fourier factor of 5/8, and with fewer diffusion directions (12 directions, total scan time ∼25 minutes). The subject was informed to move the head deliberately during the scan approximately every two minutes. The acquired Romer-EPTI data were reconstructed with and without motion correction for comparison.

### 3.3 In-vivo mesoscale dMRI at 485 μm-iso resolution at 7T

Romer-EPTI’s shortened TE, low SAR, and distortion-free imaging provide additional advantages for ultra-high field dMRI, where T_2_ is inherently shorter and field inhomogeneity is larger. To demonstrate the performance of Romer-EPTI at ultra-high field, whole-brain mesoscale in-vivo dMRI data were acquired at 485-μm iso at 7T. The key imaging parameters were: b = 1000 s/mm^2^, 40 diffusion directions, FOV = 194×186×178.5 mm^3^, matrix size of the high-resolution volume = 400×384×368, matrix size of the acquired thick-slice volume = 400×384×46, *Romer_factor_* = 8, 12 orientations (15° rotation per orientation), 3-shot EPTI encoding with a *k-t* partial Fourier factor of 5/8 (with POCS (99) for partial Fourier reconstruction), MB = 2, 50 echoes, TE_range_ = 40–110 ms, echo spacing = 1.40 ms, TR = 3 s, total acquisition time ∼75 minutes. Thick-slice EPI data were also acquired for comparison with the following parameters: FOV = 194×186×178.5 mm^3^, resolution = 0.485×0.485×3.88 mm^3^, MB = 2, R_inplane_ = 4, partial Fourier = 5/8, TE/TR = 70/3500 ms. Again, even using high accelerations (R_inplane_ = 4 and partial Fourier 5/8), EPI has a longer TE than EPTI (70 ms vs 40 ms) and a long TR due to the prolonged echo train length.

### 3.4 High b-value dMRI at b = 5000 s/mm^2^ using Romer with single-shot EPTI

High b-value dMRI often suffers from low SNR due to the strong signal decay and the long TE associated with the use of strong diffusion weightings. To demonstrate the high SNR efficiency and robustness of Romer-EPTI for high/ultra-high b-value imaging, b = 5000 s/mm^2^ dMRI data were acquired using Romer with single-shot EPTI encoding. The data were acquired at a 1.2-mm isotropic resolution with whole brain coverage on a clinical 3T scanner within 1 minute. Conventional EPI was also acquired at the same resolution within a matched 1-minute scan time. The acquisition parameters of Romer-EPTI were: FOV = 211×211×204 mm^3^, 1.2-mm isotropic resolution, matrix size of the high-resolution volume = 176×176×170, matrix size of the acquired thick-slice volume = 176×176×34, *Romer_factor_* = 5, 8 orientations (22.5° rotation per orientation), single-shot EPTI encoding with a *k-t* partial Fourier of 6/8 (with POCS), MB = 2, 66 echoes, TE_range_ = 72–133 ms, echo spacing = 0.93 ms, TR = 2.5s, number of signal averages (NSA) = 2. The acquisition parameters of EPI were: FOV = 211×211×172 mm^3^, matrix size = 176×176×144, MB = 2, R_inplane_ = 4, partial Fourier = 6/8, TE = 88 ms, TR = 10 s, NSA = 6.

### 3.5 Time-dependent diffusion experiment

As a preliminary demonstration of Romer-EPTI’s capability for imaging tissue microstructure, a time-dependent diffusion experiment was performed. Whole-brain dMRI data with 5 different diffusion times (Δ = 25, 30, 40, 50, 60 ms) were acquired using Romer-EPTI at 3T. For each diffusion time, 26 diffusion directions (12 at b = 800 s/mm^2^, 12 at b = 1600 s/mm^2^, and 2 at b = 0 s/mm^2^) were acquired, and the total acquisition time was ∼30 minutes. Other key imaging parameters were: FOV = 224×224×208 mm^3^, 2-mm isotropic resolution, matrix size of the high-resolution volume = 112×112×104, matrix size of the acquired thick-slice volume = 112×112×26, *Romer_factor_* = 4, 6 orientations (30° rotation per orientation), single-shot EPTI encoding with *k-t* partial Fourier of 6/8 (with POCS), MB = 2, 64 echoes, TE_range_ = 118–168 ms, TE was kept the same for all the diffusion times, echo spacing = 0.7 ms, and TR = 2.3 s.

### 3.6 Image reconstruction and post-processing

Image reconstruction and post-processing were performed in MATLAB. For image reconstruction, the acquired data underwent EPTI subspace reconstruction (84) with a locally low-rank constraint (regularization factor = 0.001) in BART toolbox (105,106), and then the Romer reconstruction with a Tikhonov regularization of λ = 0.025 was performed on the complex EPTI data. For image post-processing, since the reconstructed high-resolution Romer-EPTI volumes are free from distortion and dynamic distortion caused by eddy currents, only rigid registration was performed across diffusion directions using FLIRT (107,108) in FSL (109,110). FSL was then used to fit diffusion tensor imaging (DTI) (111,112) to calculate the colored Fractional Anisotropy (cFA) maps. To further improve the DTI fitting of the mesoscale dMRI data in Sections 3.1 and 3.3, local PCA (LPCA) denoising (113) was applied (note that denoising was exclusively used to calculate the mesoscale cFA maps, and no denoising was applied to any presented images (e.g., mesoscale DW images) or other results in this work). For the time-dependent dMRI experiment (Section 3.5), diffusion kurtosis imaging (DKI) (114) was fitted to the dMRI data of each diffusion time Δ using the method described in (115) to calculate the mean, radial, and axial kurtosis; a three-parameter power-law model described in (22–24,26,30) for the structural disorder was fitted to the time-dependent mean, radial, and axial kurtosis in cortical gray and white matter. Cortical gray and white matter masks were generated on a T1-weighted MPRAGE image and coregistered to the diffusion images using FreeSurfer (116–118).

## 4. Results

The point-spread-function (PSF) analysis in Figure 2A (a-b) characterizes the extent of image blurring in conventional EPI at high spatial resolution (500-μm) even with the use of a high in-plane acceleration factor of 4. This includes the T_2_/T * blurring along the phase-encoding direction due to signal decay across the readout (∼45% side-lobe amplitude, Fig. 2A-a), and the blurring in the final reconstructed volume due to the combination of encoded volumes with dynamic distortions (∼400% wider PSF, Fig. 2A-b). In contrast, by using EPTI to perform in-plane encoding of the Romer-encoded volumes, these sources of blurring can be eliminated (Figure 2A-c). To characterize the effective resolution (along the slice-readout directions) and SNR of the Romer reconstruction, Figure 2A-d shows the PSF-error versus 1/SNR plots (L-curve) under different regularization parameters. As can be seen, at a good trade-off point between PSF error and SNR (λ = 0.025), a Romer factor of 8 in acquisition can achieve a high SNR efficiency gain of 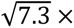 (close to the theoretical SNR gain of √8 ×), while achieving sharp resolution along slice-readout dimensions with mild spatial blurring (∼14% side-lobe amplitude) at a mesoscale resolution of 500 μm. This, combined with SMS and optimized EPTI readout, achieves an overall SNR efficiency gain of 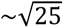 (see calculation in section 2.2). Figure 2B shows the high isotropic spatial resolution reconstructed by Romer (middle, 0.5-mm iso) compared to the directly acquired high in-plane resolution EPTI image as a reference (right, 0.5×0.5 mm^2^ in-plane). The Romer reconstructed image shows sharp structures with no noticeable blurring, demonstrating its effectiveness in reconstructing high isotropic spatial resolutions. The low-resolution view of one of the thick-slice volumes is also shown (left, 4×0.5 mm^2^) for comparison.

The whole-brain distortion-free in-vivo mesoscale dMRI data acquired by Romer-EPTI at 500-μm iso on a 3T clinical scanner are shown in Fig. 3-5. The mean DWI images (Fig. 3A) show exceptional isotropic resolution in three orthogonal views, revealing fine-scale structures such as the intricate layered cortical folding in the hippocampus and cerebellum, as well as detailed subcortical structures including deep brain basal ganglia such as the caudate and putamen, and brain stem nuclei. At this mesoscale spatial resolution, these DWI images present superior quality with high SNR and sharp spatial resolution, which are also free from distortion and slab-boundary artifacts. Notably, the deep brain regions, which typically suffer from severe distortion and low SNR in conventional acquisition, show high SNR and anatomical integrity in the Romer-EPTI images. The single-direction DWI images are also presented (Fig. 3B).

**Figure 3.**
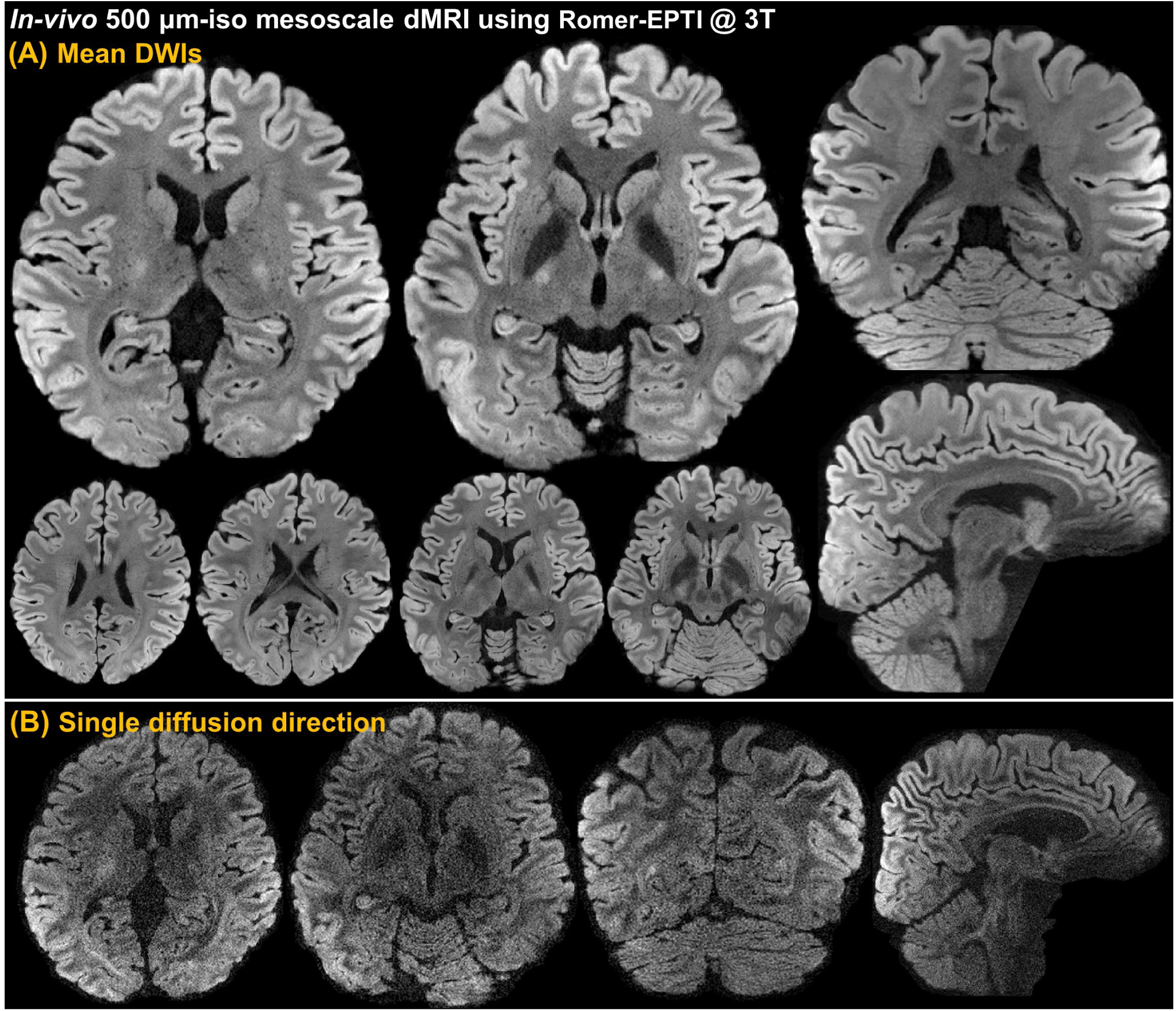
In-vivo whole-brain distortion-free 500 μm-isotropic mesoscale diffusion data acquired by Romer-EPTI on a clinical 3T scanner. (**A**) Mean diffusion-weighted images (DWIs) of different axial, coronal, and sagittal views (b = 1000 s/mm^2^). At this mesoscale spatial resolution, the diffusion images exhibit superior image quality characterized by their high SNR, distortion-free, no slab boundary artifacts, and sharp isotropic resolution. The mesoscale resolution reveals exquisite fine-scale structures such as the intricate layered cortical folding in the hippocampus and cerebellum, as well as detailed subcortical structures including deep brain basal ganglia such as the caudate and putamen, and brain stem nuclei. The deep brain regions, which typically suffer from severe distortion and low SNR in conventional acquisition, show high SNR and quality in the Romer-EPTI image. (**B**) The single-direction DWIs of the Romer-EPTI data, which are also free from distortion and slab boundary striping artifacts.

**Figure 4.**
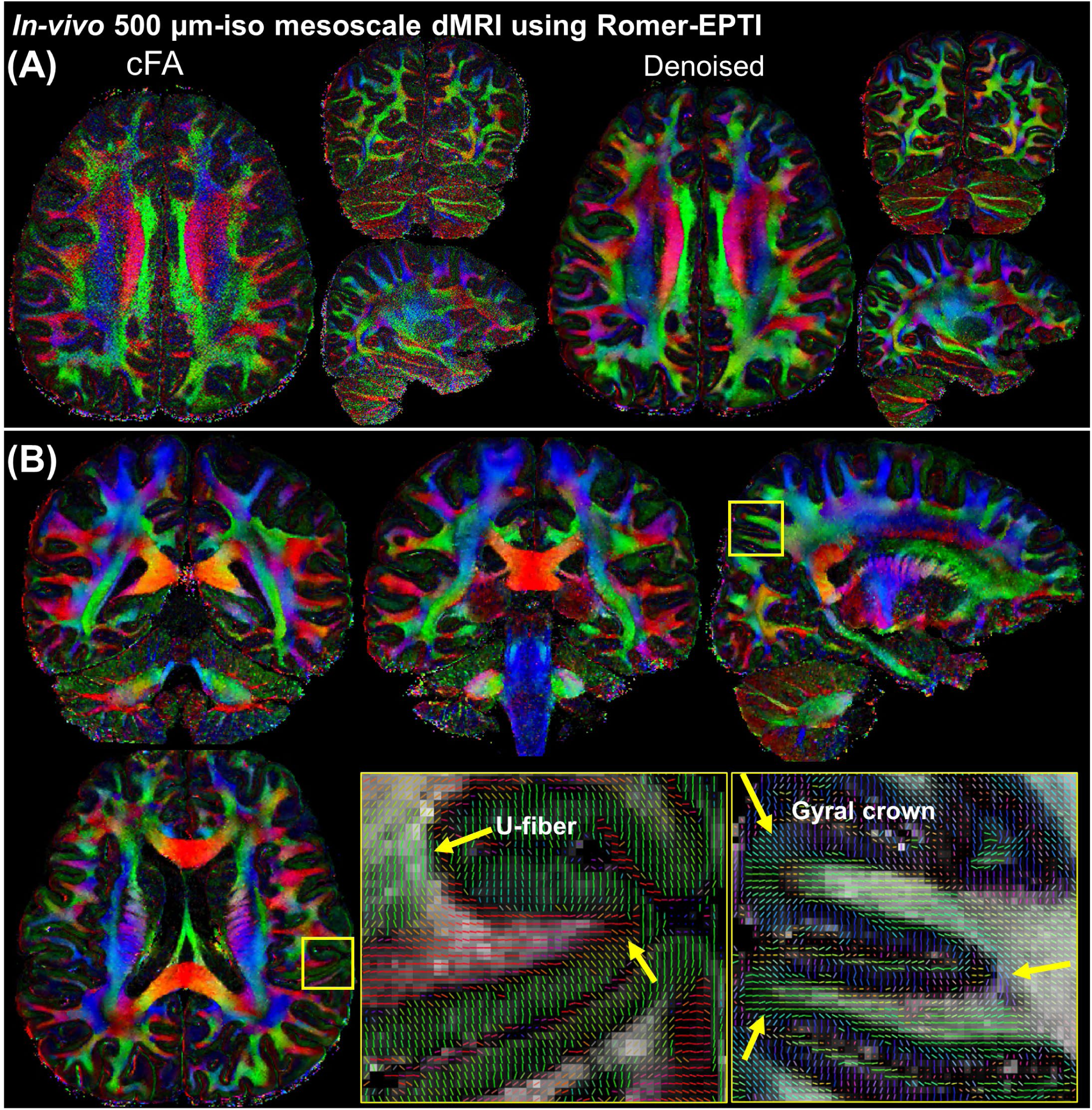
Colored-FA (cFA) maps calculated from the 500 μm-iso mesoscale dMRI Romer-EPTI data at 3T. (**A**) The mesoscale cFA maps acquired by Romer-EPTI without and with LPCA denoising. The high spatial resolution with sharp details is well-preserved in the denoised cFAs while the noise level is further reduced. (**B**) mesoscale cFA maps reveal fine-scale structures and connectivity not only in white matter but also within the thin cortical gray matter. The zoomed-in views of the diffusion tensors highlight the resolved small and rapidly turning U-fibers at the gray-white matter boundary and gyral crown within the thin cortex (yellow arrows). Note that denoising was exclusively used to compute the mesoscale cFA maps, and no denoising was applied to any presented images (e.g., mesoscale DW images) or other results in this study.

**Figure 5.**
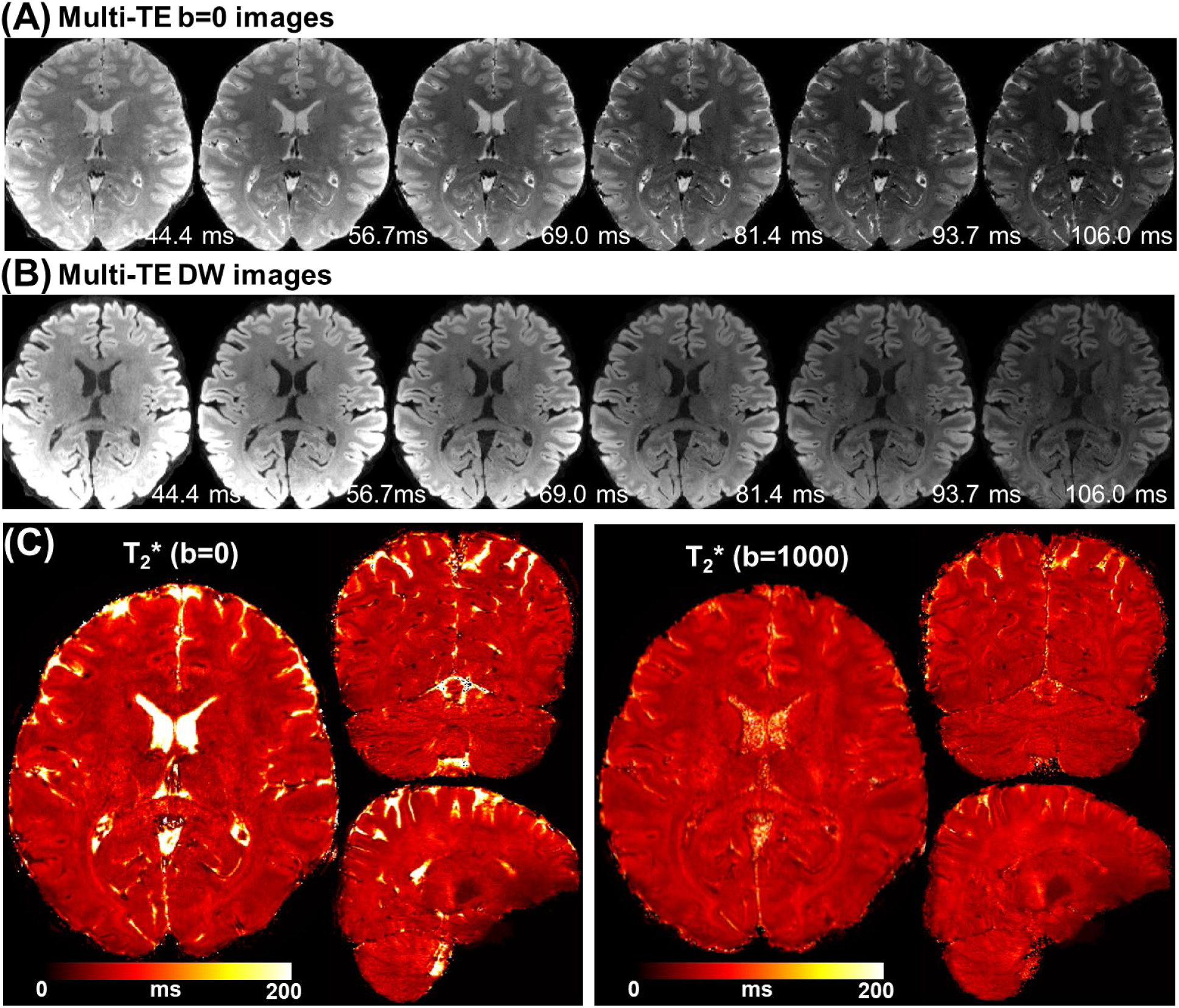
Multi-TE dMRI data at 500 μm-isotropic resolution simultaneously acquired by Romer-EPTI. (**A**) Multi-TE b=0 s/mm^2^ images of 6 averaged echo groups from a total of 48 echoes. The effective TE of each echo group is shown. (**B**) Multi-TE DWI images (b = 1000 s/mm^2^) of the same 6 echo groups. (**C**) Fitted T_2_* map using the multi-TE b=0 and DWI images presented in 3 orthogonal views. The DWI T_2_* maps exhibit overall lower T_2_* values than the b=0 as expected because of the diffusion weighting’s suppression effect on the free water compartment.

The calculated colored-FA maps from this data are shown in Figure 4. At such ultra-high resolution (∼0.1 mm^3^), the cFA maps still exhibit some noise, and after LPCA denoising, the cFA maps show increased SNR with preserved spatial resolution (Fig. 4A). More views of the cFA maps are presented in Fig. 4B, exhibiting high quality and SNR. In addition to the sharp fibers in the white matter, achieving such high mesoscale resolution allows the investigation of rapidly turning fibers at the gray-white matter boundaries (e.g., U-fibers) and within the thin cortical gray matter (e.g., gyral structures), as shown by the zoomed-in tensor images. Note that denoising was exclusively used to compute the mesoscale cFA maps, and no denoising was applied to any presented images (e.g., mesoscale DW images) or other results in this study.

Figure 5 shows the 500-μm b=0 and diffusion-weighted images with simultaneously acquired multi-TEs. Consistent high image quality and spatial resolution were observed across different TEs. The fitted T_2_* maps using the b=0 s/mm^2^ and b=1000 s/mm^2^ data are also shown. An overall lower T_2_* value was observed in the b=1000 s/mm^2^ T * maps compared to the b=0 s/mm^2^ maps as expected due to the suppression of the free water compartment (long T_2_/T_2_*) by the diffusion gradients.

Figure 6 shows the results of the motion experiment, which compares the 500-μm isotropic Romer-EPTI data reconstructed without and with the motion-aware reconstruction. The subject exhibited translation and rotation movements up to ∼±6 mm/degrees (estimated motion shown in Supporting Information Figure S2). Without the motion-aware reconstruction, severe image blurring and artifacts appear in both the b=0 s/mm^2^ and DWI images. In contrast, using the proposed motion-aware Romer reconstruction, high-resolution images are reconstructed with successfully restored sharp details as highlighted by the zoomed-in views. This shows the capability of Romer-EPTI to produce high-quality dMRI data at high spatial resolution even in the presence of substantial subject motion.

**Figure 6.**
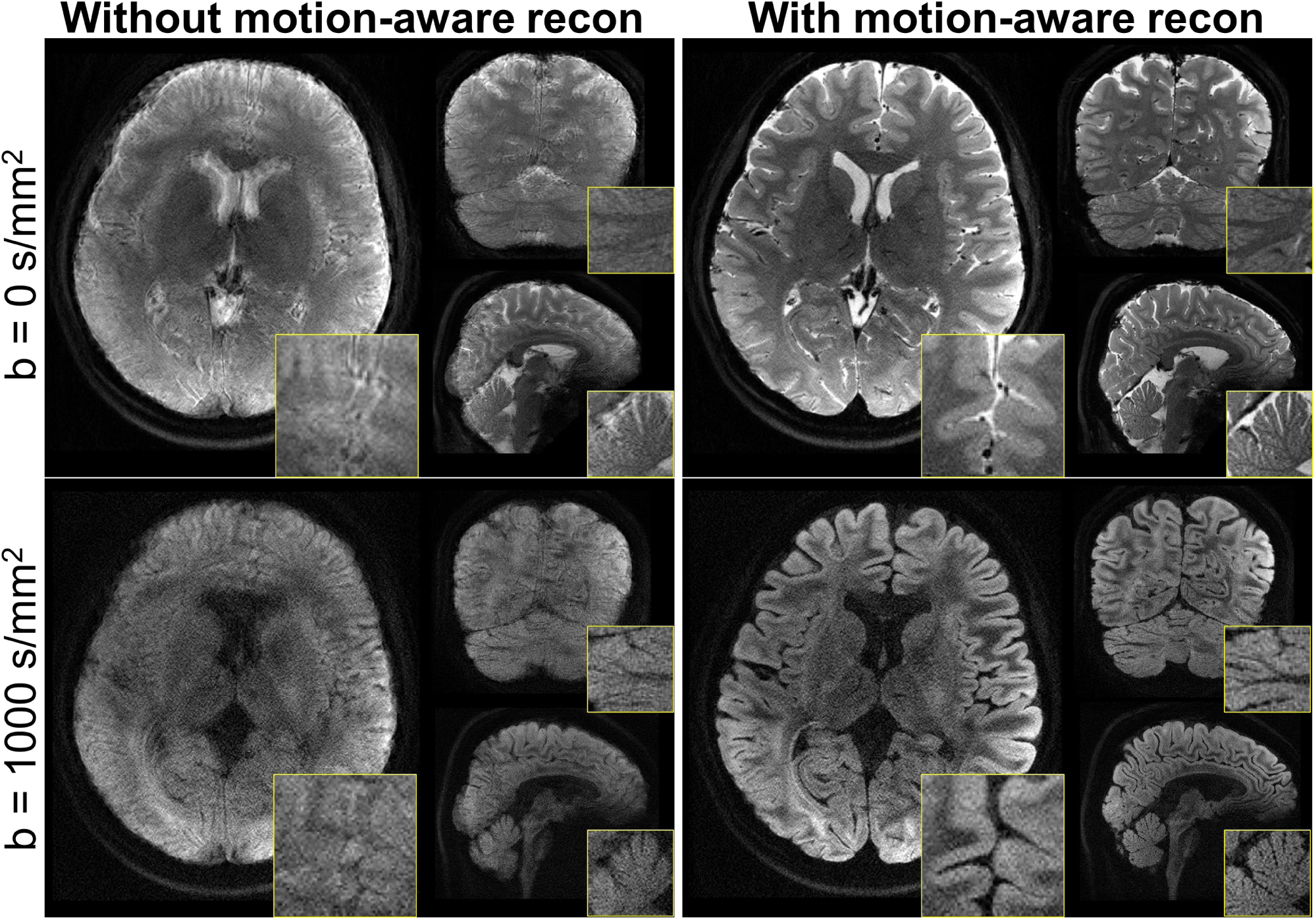
Results of in-vivo motion experiment comparing non-DW and DWI Romer-EPTI images without and with the motion-aware reconstruction at mesoscale 500 μm-iso at 3T. The subject exhibited translation and rotation movements up to ∼±6 mm/degrees (estimated motion shown in Supporting Information Figure S2). Without the motion-aware reconstruction, severe image blurring and artifacts appear in both the b=0 s/mm^2^ and DWI images. In contrast, the proposed motion-aware reconstruction cleans up these blurring artifacts and provides sharp high-resolution images with high image quality, as can be seen in the zoomed-in images, demonstrating the high motion robustness of Romer-EPTI.

At 7T, Romer-EPTI enables high-quality, distortion-free whole-brain in-vivo mesoscale dMRI (Fig. 7). High-resolution dMRI on 7T poses greater challenges than 3T due to stronger field inhomogeneity and faster T_2_/T_2_* decay. By shortening TE (e.g., 40 ms vs 70 ms), Romer-EPTI can better take advantage of the SNR gain offered by the increased field strength with less offsets from the shorter T_2_* (see improved SNR observed in 7T images compared to 3T in Supporting Information Figure S3). Benefiting from the higher SNR at 7T, we further increased the spatial resolution to 485-μm isotropic. The acquired mean DWI images (Fig. 7A) show high SNR and high image quality without distortion or striping artifacts. The 7T data exhibit a stronger gray-white matter contrast. The mesoscale spatial resolution reveals exquisite fine-scale structures, such as i) the thin edges of the putamen and white matter tracts, ii) the layered cortical folding in the hippocampus, and iii) the gray matter bridges spanning the internal capsule, highlighting the high spatial resolution and accuracy of this data. Image intensity inhomogeneities due to B_1_ inhomogeneity are observed, particularly at the bottom part of the brain. The colored-FA maps shown in Fig. 7B also demonstrate high SNR at this mesoscale resolution.

**Figure 7.**
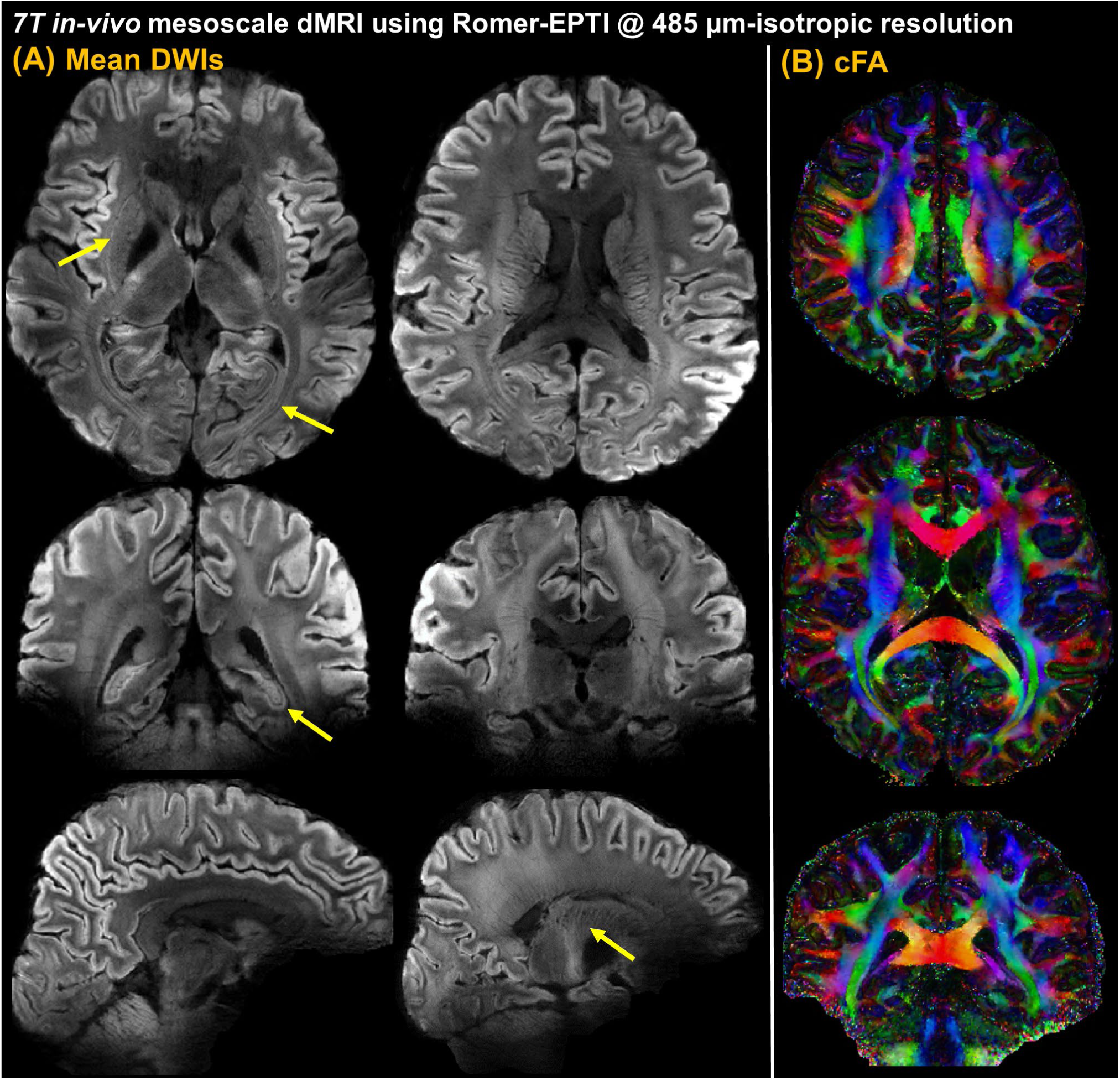
In-vivo whole-brain distortion-free 485 μm-isotropic mesoscale dMRI data on a clinical 7T scanner acquired by Romer-EPTI. (**A**) The mean DWIs exhibit high SNR and sharp spatial resolution, revealing exquisite fine-scale structures highlighted by the yellow arrows, including the thin edges of the putamen and white matter tracts (axial view), the layered cortical folding in the hippocampus (coronal view), and the gray matter bridges spanning the internal capsule (sagittal view), highlighting the minimal blurring and high spatial accuracy of this data. There are no distortion or striping artifacts observed in this mesoscale data, and the gray-white matter contrast at 7T is much stronger than 3T. Image intensity inhomogeneities resulting from strong B_1_ inhomogeneity at 7T are observed, particularly at the bottom part of the brain, which can be mitigated through parallel transmission. (**B**) The colored-FA maps of the 7T mesoscale data.

Figure 8 evaluates the image quality of the Romer-EPTI data at mesoscale resolutions (500-μm isotropic at 3T, 485-μm isotropic at 7T), and compares it with conventional SMS-EPI with the same in-plane resolution but thicker slices (0.5×0.5×4 mm^3^ at 3T and 0.485×0.485×3.88 mm^3^ at 7T). In both the 3T and 7T images, conventional EPI images show severe distortion artifacts (indicated by red arrows) and blurring even with a high in-plane acceleration factor of 4, along with ghosting artifacts (orange arrows). In contrast, Romer-EPTI provides distortion-free and sharp images with high SNR. Note that thick slices were used in EPI (see the coronal views) to boost its SNR, roughly matching that of Romer-EPTI without significantly extending the scan time (to achieve the same isotropic resolution at a similar SNR, EPI would take >30 minutes), for a fair image quality comparison for EPI.

**Figure 8.**
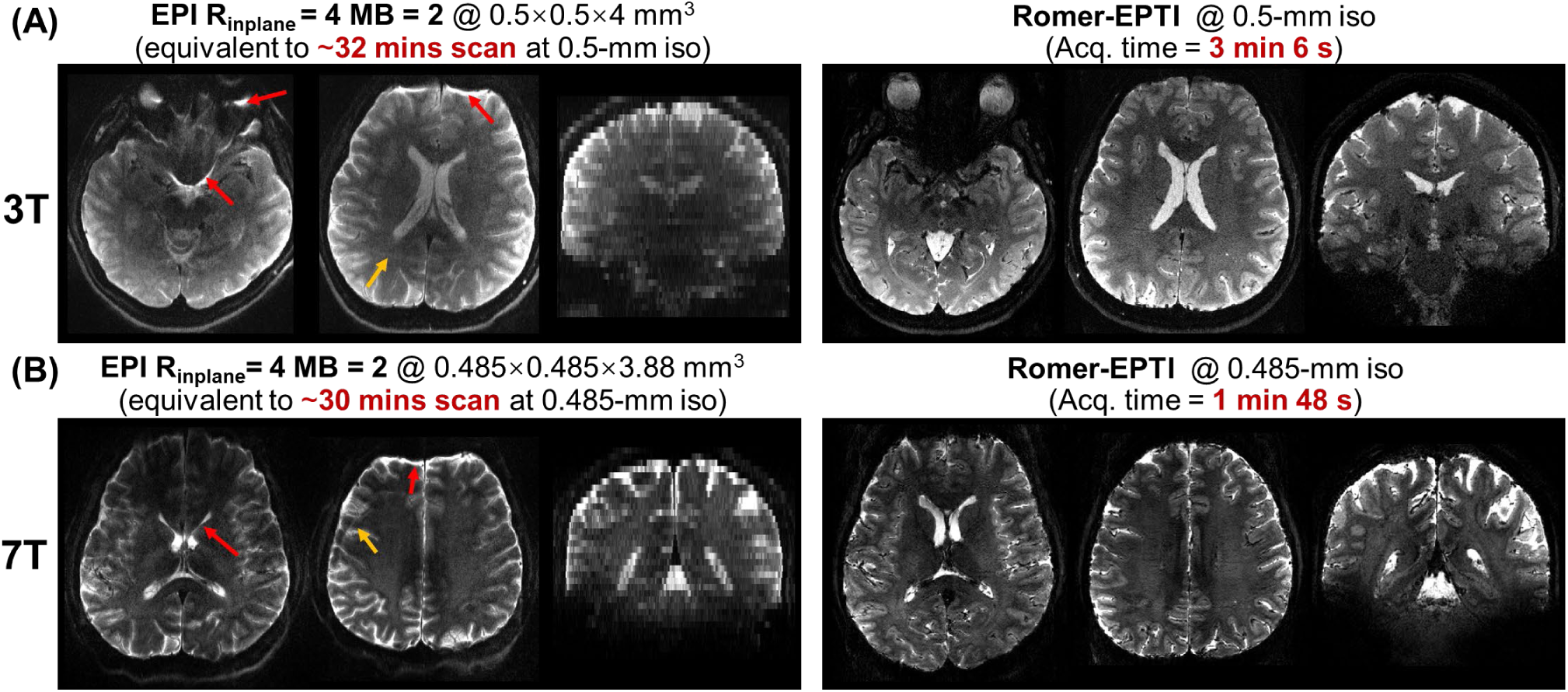
Image quality comparison at 3T **(A)** and 7T **(B)** between Romer-EPTI at isotropic mesoscale resolution and conventional SMS-EPI at the same in-plane resolution. In both the 3T and 7T images, conventional EPI images show severe distortion artifacts (red arrows) and blurring even with a high in-plane acceleration factor of 4, along with ghosting artifacts (orange arrows). In contrast, Romer-EPTI provides distortion-free and sharp images with high SNR. Note that thick slices were used in EPI (see coronal views) to boost its SNR, roughly matching that of Romer-EPTI without significantly extending the scan time (to achieve the same isotropic resolution at a similar SNR, EPI would take >30 minutes), for a fair image quality comparison for EPI.

The whole-brain high b-value imaging at b=5000 s/mm^2^ and 1.2-mm isotropic resolution are shown in Fig. 9. As can be seen, Romer-EPTI provides significantly higher SNR compared to the SMS-EPI data acquired within a matched 1-minute scan time. The use of single-shot EPTI encoding ensures high robustness to strong phase variations in high b-value imaging. Figure 10 shows the results of the time-dependent diffusion experiment. The mean, radial, and axial kurtosis maps derived from the whole-brain Romer-EPTI data acquired at 2-mm isotropic resolution at five different diffusion times are presented. The calculated mean, radial and axial kurtosis reveal clear time-dependency in both cortical gray matter and white matter (p-value <0.05).

**Figure 9.**
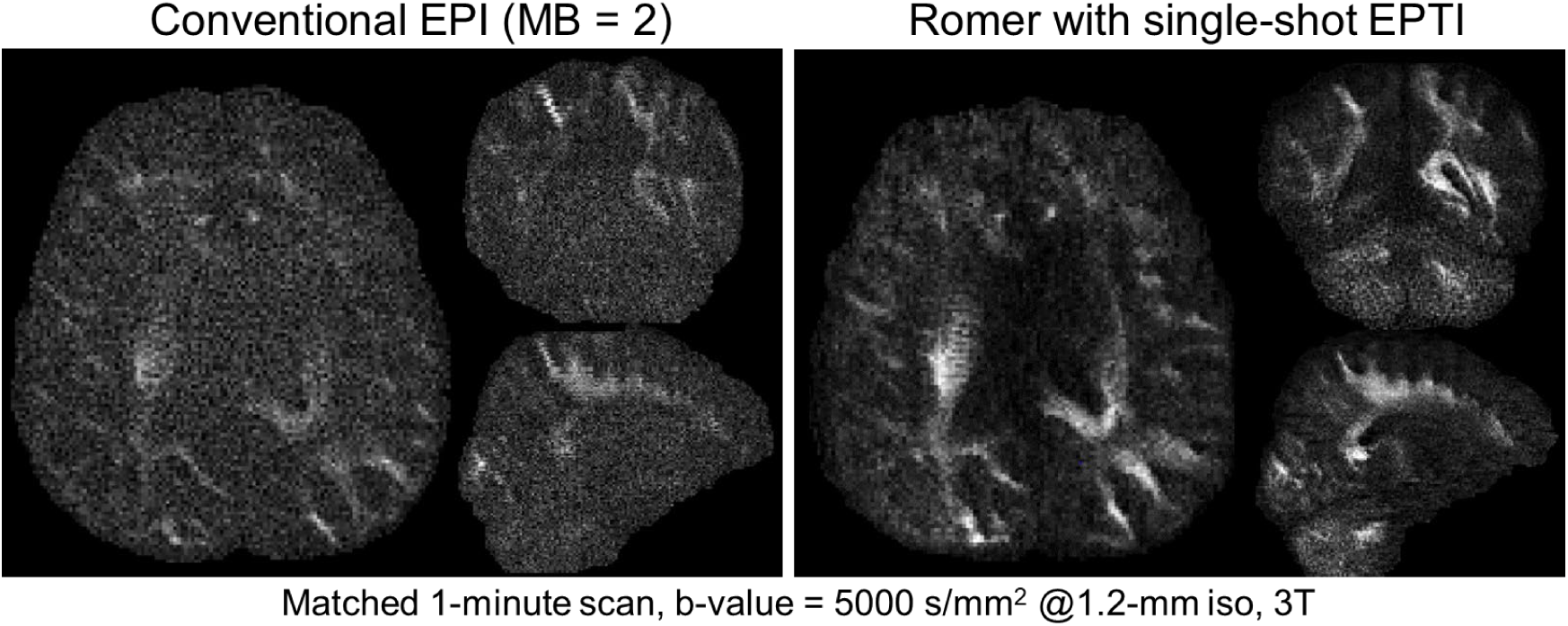
High b-value dMRI results at b = 5000 s/mm^2^ and 1.2-mm isotropic resolution using conventional SMS-EPI and Romer-EPTI. In a matched 1-minute scan time, Romer-EPTI provides significantly higher SNR compared to the conventional SMS-EPI data, demonstrating the SNR gain and utility of Romer-EPTI for high b-value dMRI. The single-shot EPTI encoding ensures high robustness to strong phase variations in high b-value scans.

**Figure 10.**
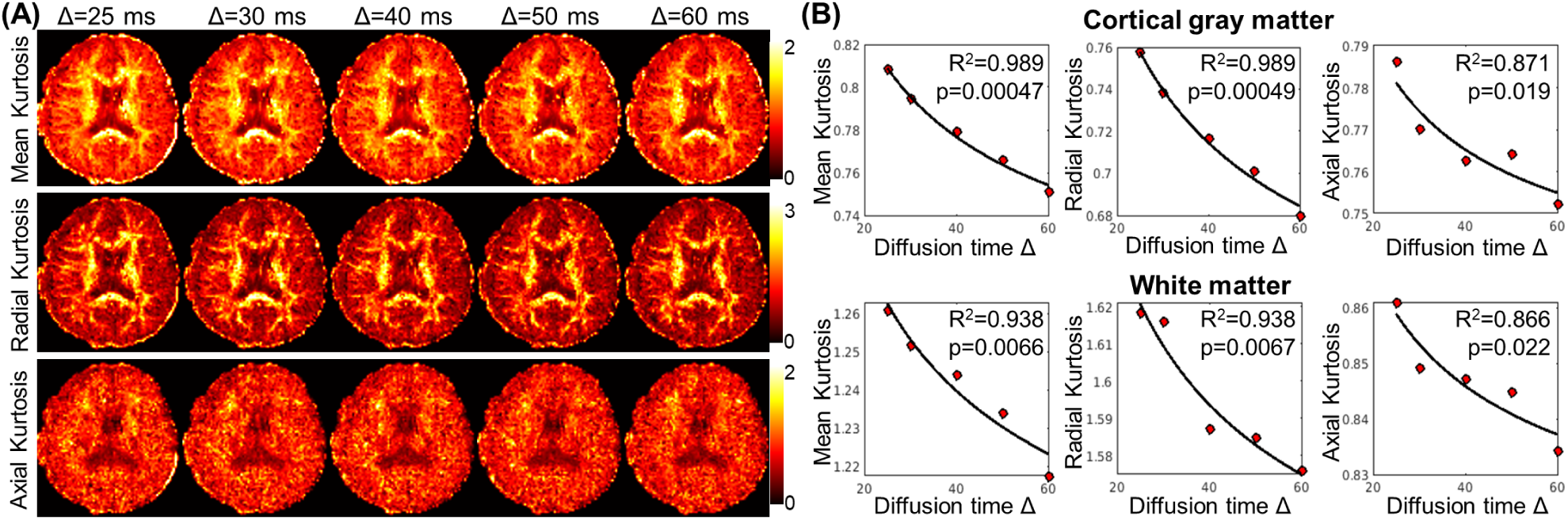
Results of the time-dependent diffusion experiment, where Romer-EPTI data at 2-mm isotropic resolution with whole-brain coverage at five different diffusion times (Δ=25, 30, 40, 50, 60 ms) were acquired (each diffusion time acquired in ∼6 minutes with 26 diffusion directions). (**A**) Mean, radial, and axial kurtosis maps derived from the data at the five diffusion times. (**B**) Mean, radial, and axial kurtosis in cortical gray matter and white matter plotted as a function of diffusion time, where clear time-dependence is observed (p<0.05).

## 5. Discussion and Conclusions

This study aims to overcome the major challenges in diffusion MRI acquisition, including low SNR, poor image quality, and susceptibility to motion artifacts—especially prominent when aiming for higher spatial resolutions or increased b-values. To achieve this, we introduce a novel Romer-EPTI technique, which integrates an SNR-efficient rotating-view motion-robust super-resolution technique with a distortion/blurring-free self-navigated EPTI encoding. We demonstrated that Romer-EPTI provides significant gain in SNR efficiency (e.g., 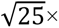), improved image quality free from distortion, T_2_/T_2_* blurring, and slab-boundary artifacts, the ability to resolve sharp spatial resolution, as well as high robustness to various sources of motion and additional multi-echo information. These are all achieved without introducing extra SAR or *B*_0_/*B*_1_ susceptibility. Using Romer-EPTI, we successfully acquired whole-brain mesoscale in-vivo dMRI at both 3T and 7T, and to our knowledge, this is the first study that has achieved this. Furthermore, Romer-EPTI’s efficacy for efficient and robust microstructure imaging has also been demonstrated through high b-value and time-dependent diffusion experiments.

The whole-brain mesoscale in-vivo dMRI data acquired at 3T and 7T demonstrate the significant SNR gain and superior image quality provided by Romer-EPTI. The spatial resolution achieved in this study (∼0.12 mm^3^) is ∼27-fold higher than the standard dMRI acquisition at 1.5 mm isotropic resolution (∼3.3 mm^3^), and ∼4-fold higher than the high-performance 14-hour submillimeter dataset (0.44 mm^3^) we have recently published (8), providing much more detailed structural information to aid in future studies (119). Microstructure imaging demonstrated here, such as high-b value (15–19,27) and time-dependent imaging (22–26,30), represents another vital application in dMRI. Using conventional acquisition, even at moderate/standard spatial resolution, these microstructure imaging applications can still be SNR-starved due to the strong signal decay and long TE associated with high b values. In addition, the diffusion model fitting can be sensitive to data’s SNR levels and may require extended scan times (24,26). The successful measurement of time-dependent kurtosis in an efficient acquisition using Romer-EPTI (each diffusion time acquired in 6 mins with 26 diffusion directions and whole brain coverage) preliminarily validates its efficacy and potential to serve as an efficient acquisition tool for dMRI-based microstructure imaging.

Romer-EPTI’s high SNR efficiency significantly reduces the scan time for mesoscale dMRI and ultra-high b-value imaging. For example, the conventional 2D EPI acquisition would require >30 hours to reach the same SNR level at the same mesoscale resolution of ∼485 μm-iso achieved in our 75-minute Romer-EPTI data. The high SNR gain of Romer-EPTI (e.g., 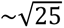 folds) results from the combination of the simultaneous multi-thick-slice Romer acquisition and the efficient EPTI readout with minimized TE and optimized readout length. Further increasing the Romer factor can provide higher SNR efficiency, but careful consideration of two factors is essential. First, a higher Romer factor leads to a thicker slice thickness, and the capability of the thicker-slice volume to accurately estimate motion and physiological phase should be considered. Second, although a higher Romer factor will reduce the volume TR, pushing the reduction of volume TR beyond a certain point (e.g., < 2 s) may yield less gain in SNR efficiency for spin-echo-based diffusion sequences, as the magnetization cannot effectively recover within each TR. Moving to 7T, Romer-EPTI addresses unique challenges compared to 3T. At 7T, in conventional dMRI acquisition, where long EPI readouts and extended TE are employed, the anticipated SNR gain from increased field strength is offset by the inherently shorter T_2_ at 7T. In contrast, the minimized TE (e.g., 40 ms vs 70 ms) achieved by Romer-EPTI can better preserve the SNR gain from the increased field strength with less offset from the shorter T_2_*. The distortion-free nature of Romer-EPTI also addresses the stronger *B*_0_ inhomogeneity at 7T. The *B*_1_ inhomogeneity is another common problem for spin-echo dMRI sequences at 7T, and future work will integrate parallel transmission and pTx pulses (120–122) with Romer-EPTI to address this.

Advanced denoising techniques and hardware can be combined with Romer-EPTI to further enhance SNR. For instance, our demonstration preliminarily shows that applying denoising (LPCA) (113) to Romer-EPTI data effectively enhances the SNR of the FA maps. Exploring new denoising methods (123–127) with finely tuned parameters tailored to Romer-EPTI’s data and noise characteristics, or employing machine learning-based denoising techniques (128,129), holds promise for further improving SNR. Moreover, all data presented in this study were acquired on clinically accessible 3T and 7T scanners using commercially available coils. Future work will leverage high-performance MRI systems, such as Connectome 2.0 (11), MAGNUS (12), and ‘Impulse’ 7T (130), along with custom-built high-channel receive arrays (131) for further improvement in achievable spatial resolution, acquisition efficiency, and attainable b-values for in-vivo dMRI.

In addition to SNR, the ability of Romer-EPTI to effectively address distortion, blurring, and motion-related artifacts is critical in achieving mesoscale and high b-value dMRI, as these problems become more prominent when increasing spatial resolution and/or b-values. Romer-EPTI’s robustness to motion and phase variation benefits from its self-navigation nature. Each Romer-encoded thick-slice volume is reconstructed independently with high SNR and can be used to estimate bulk motion and phase variations. This makes the motion correction problems more tractable, as it decouples motion correction from unaliasing using parallel imaging and *k*-space reconstruction (e.g., multi-slab imaging), or from interference with RF encodings that introduce encoding-related contrast differences and phase changes that complicate motion and phase estimation (e.g., gSlider) (43). Another key element contributing to Romer’s high motion robustness is the utilization of distortion-free EPTI readout, which effectively eliminates the dynamic distortions/geometric inconsistencies across the thick-slice volumes that can be induced by subject motion. Such an effect is challenging to estimate and correct and has therefore often been overlooked, but can result in detrimental spatial blurring in super-resolution reconstruction (Fig. 2A-b). In the context of mesoscale dMRI, multi-shot EPTI was employed for the in-plane encoding of Romer to ensure favorable reconstruction conditioning for achieving ultra-high spatial resolution, and distortion/blurring-free mesoscale in-plane resolution was achieved using 3-shot EPTI. The EPTI encoding was designed with efficient self-navigation that can effectively correct for shot-to-shot phase variations. Since the motion between different Romer-encoded volumes has been modeled, the motion sensitivity timeframe of Romer-EPTI at mesoscale spatial resolution is short (e.g., volume TR of 9 s using 3 shots at 485 μm) compared to EPI (e.g., volume TR of ∼30 s) or other thick-slab/slice acquisitions (e.g., >1 minute if at the same resolution and coverage (39,77)). The self-navigator of EPTI can be used to further reduce the motion sensitivity timeframe of mesoscale dMRI down to 2-3 s, and future work will exploit this for intra-encoded-volume motion correction (43,132). It is also worth noting that, even without this intra-encoded-volume motion correction and data corruption occurs for certain thick-slice volumes, they can be simply removed before Romer reconstruction without severely degrading the quality of the final reconstructed high-resolution volume, benefiting from the oversampled center with data redundancy of the rotating-view encoding. In the context of high b-value dMRI and microstructure imaging, where the strong diffusion gradient makes these applications sensitive to shot-to-shot phase variations and data corruption, single-shot EPTI was employed for in-plane encoding that can achieve high b-value dMRI at sufficiently high spatial resolutions (e.g., 1-2 mm) in a single shot acquisition, avoiding the intra-volume shot-to-shot phase variations and motion issues. This already achieved a motion sensitivity timeframe down to 2-3 s, making Romer-EPTI a highly efficient and motion-robust acquisition for high b-value dMRI and microstructure imaging.

Other optimizations in Romer-EPTI are also important in achieving high image quality and accurate reconstruction with minimal bias. Specifically, the acquisition scheme in Romer-EPTI was designed to avoid accumulative spin history effect and spatially non-uniform signal recovery, therefore providing images free from common striping artifacts. In addition, given that Romer encoding does not depend on RF encoding to resolve spatial resolution, it is inherently less susceptible to *B*_0_/*B*_1_ imperfections. For accurate super-resolution reconstruction, realistic slice profile modeling in Romer avoids residual biases (Supporting Information Figure S1B). Factors affecting the slice profile, such as *B*_0_ and *B*_1_ inhomogeneity, can also be modeled for each voxel to further improve the accuracy in the future. Moreover, the complex-valued reconstruction was designed to enable real-value dMRI (95), which effectively reduces the noise floor (Supporting Information Figure S1A).

In addition to high-performance dMRI for neuroscientific applications, Romer-EPTI also holds great promise for clinical applications, where it can increase spatial resolution, enable efficient microstructure measurements, and enhance image quality with matched anatomy integrity as distortion-free anatomical scans. Based on the demonstrated mesoscale dMRI at 485 μm-iso, whole-brain dMRI at 750 μm-iso can be acquired in just ∼7-8 minutes with the same SNR level. Romer-EPTI can also be further accelerated. For instance, the current Romer reconstruction is performed independently for each diffusion encoding volume, but the rotating-view encoding of Romer makes it well-suited for undersampled reconstruction and acceleration methods, such as joint diffusion reconstruction (133–135), which can further reduce the scan time and enhance the efficiency of Romer-EPTI.

## Acknowledgements

This work was supported by the NIH NINDS *BRAIN Initiative* (U24NS129893), NIA (K99AG083056), NIBIB (P41EB030006, U01EB026996, R01EB019437), NIDCR OD (DP5OD031854), and by the MGH/HST Athinoula A. Martinos Center for Biomedical Imaging, and was made possible by the resources provided by NIH Shared Instrumentation Grants S10OD023637.

---

**Figure S1.**
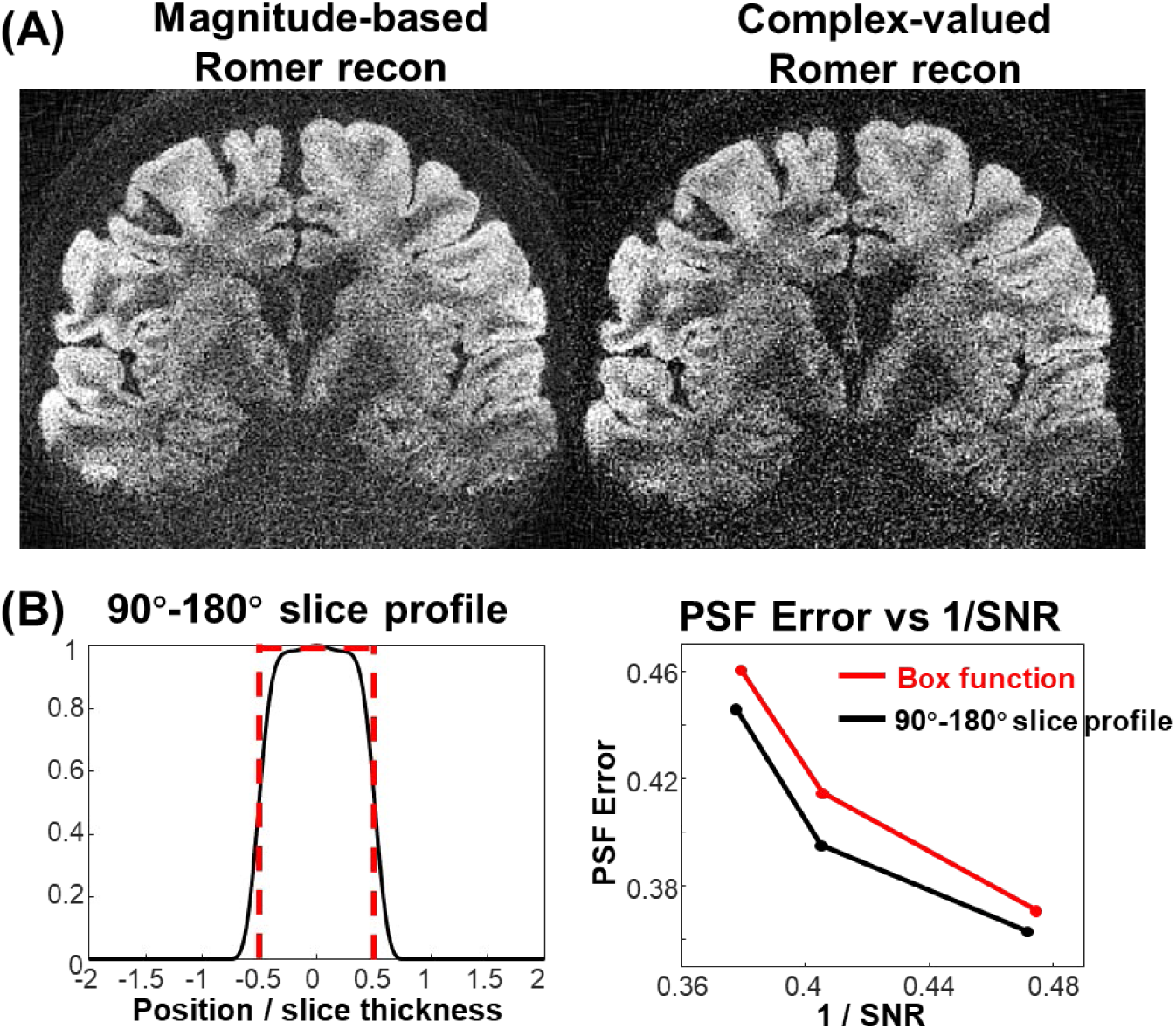
(**A**) Comparison of the magnitude-data-based and the complex-valued Romer reconstruction results. The noise is effectively reduced using the proposed complex-valued Romer reconstruction compared to the magnitude-based reconstruction. Note that the low-frequency phase differences between the thick-slice volumes are removed in the complex-valued reconstruction. (**B**) The realistic slice profile calculated based on the 90°- 180° RF pulses. By modeling the realistic slice profile in Romer reconstruction, residual PSF error can be removed compared to using a simplified Box function.

**Figure S2.**
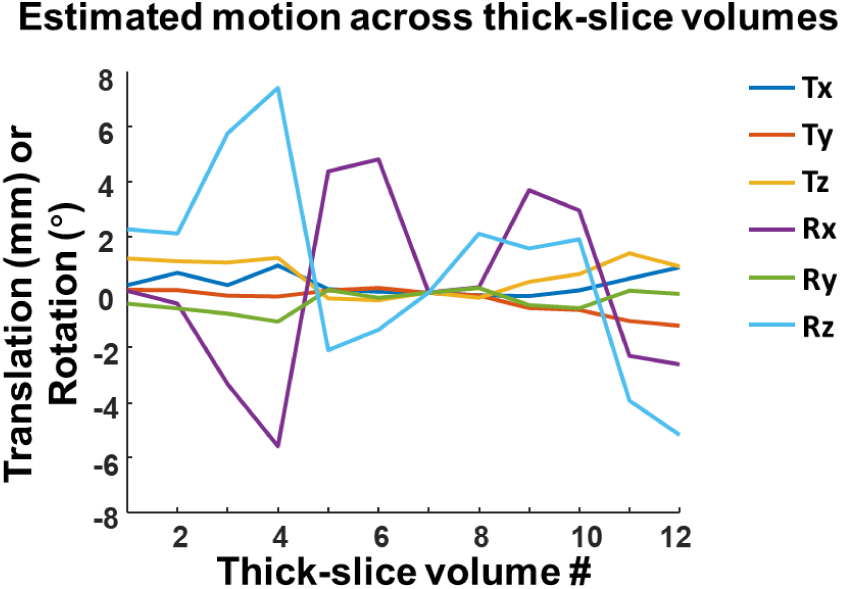
The estimated rigid motion parameters between the encoded thick-slice volumes during the in-vivo motion experiment.

**Figure S3.**
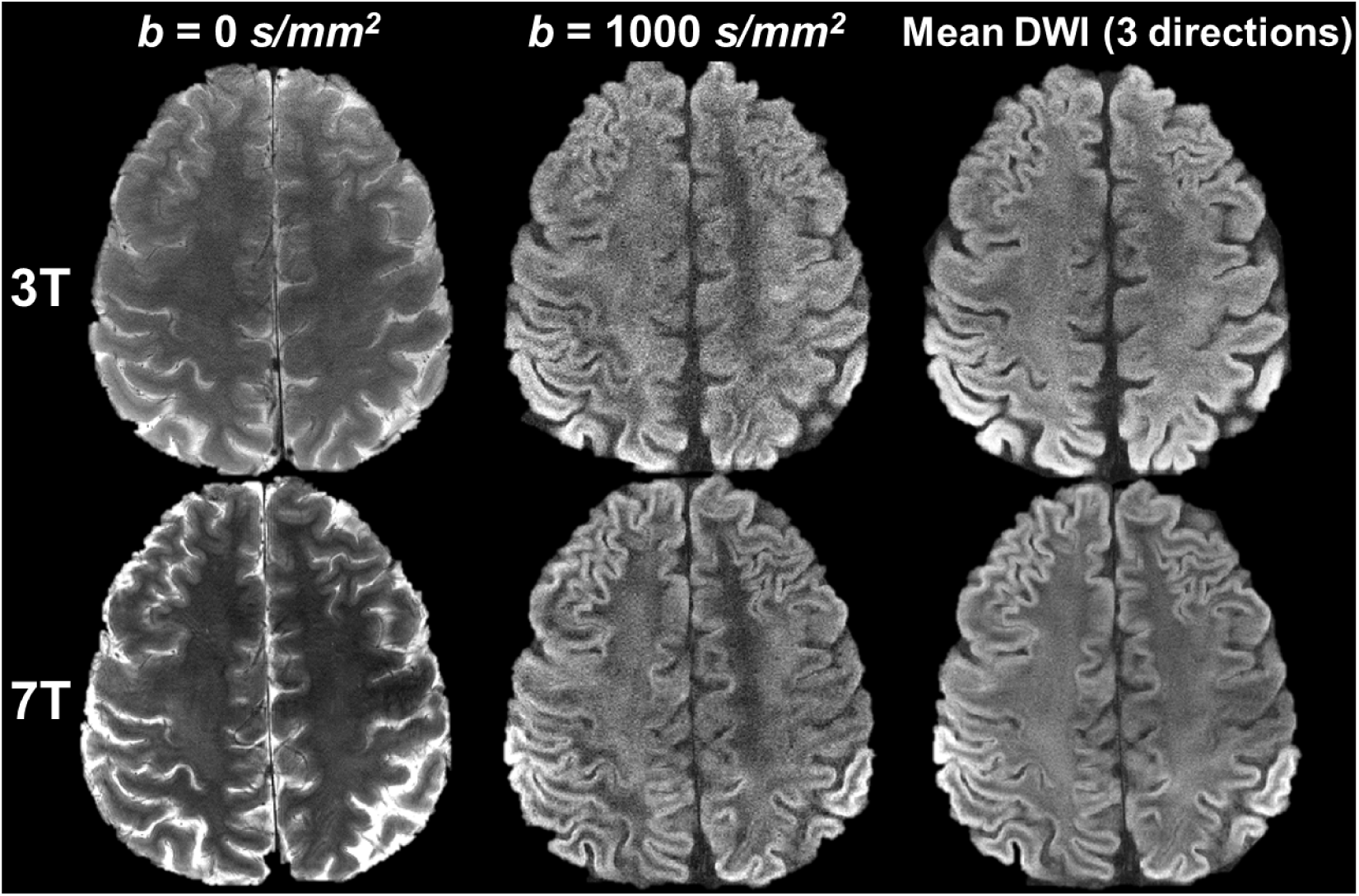
Comparison of EPTI images acquired at 3T and 7T. The b=0 s/mm^2^ images, b=1000 s/mm^2^ images, and the mean DWIs of 3 diffusion directions are shown. The 7T images show higher SNR compared to 3T. The SNR improvement resulting from EPTI’s minimized TE is especially beneficial for 7T dMRI where the T_2_ signal decay is inherently faster. During the comparison, both 3T and 7T data were acquired with the same image resolution at 0.5×0.5×4 mm^3^, and the same spin-echo TE of 39 ms.

